# Active site plasticity and possible modes of chemical inhibition of the human DNA deaminase APOBEC3B

**DOI:** 10.1101/513366

**Authors:** Ke Shi, Özlem Demir, Michael A. Carpenter, Surajit Banerjee, Daniel A. Harki, Rommie E. Amaro, Reuben S. Harris, Hideki Aihara

## Abstract

The single-stranded DNA cytosine deaminase APOBEC3B (A3B) functions in innate immunity against viruses, but is also strongly implicated in eliciting mutations in cancer genomes. Because of the critical role of A3B in promoting virus and tumor evolution, small molecule inhibitors are desirable. However, there is no reported structure for any of the APOBEC3-family enzymes in complex with a small molecule bound in the active site, which hampers the development of small molecules targeting A3B. Here we report high-resolution structures of an active A3B catalytic domain chimera with loop 7 residues exchanged with those from the corresponding region of APOBEC3G (A3G). The structures reveal novel open conformations lacking the catalytically essential zinc ion, with the highly conserved active site residues extensively rearranged. These inactive conformations are stabilized by 2-pyrimidone or an iodide ion bound in the active site. Molecular dynamics simulations corroborate the remarkable plasticity of the engineered active site and identify key interactions that stabilize the native A3B active site. These data provide insights into A3B active site dynamics and suggest possible modes of its inhibition by small molecules, which would aid in rational design of selective A3B inhibitors for constraining virus and tumor evolution.

## Introduction

The hallmark activity of the APOBEC family of enzymes is catalyzing a zinc ion-mediated hydrolytic deamination of 2′-deoxycytidine to 2′-deoxyuridine in single-stranded (ss)DNA (reviewed by refs.^1,2^). In addition to AID and APOBEC1, human cells have the potential to express up to seven different APOBEC3 (A3) enzymes (A3A-D, A3F-H). These enzymes have overlapping functions in providing innate immunity to viral infections with, for instance, A3D/F/G/H capable of restricting the replication of the retrovirus HIV-1. Although best characterized for roles in restricting retrovirus and retrotransposon replication (with obligate ssDNA intermediates), several family members have also been implicated in the restriction of small and large double-stranded (ds)DNA tumor viruses. For instance, A3B and A3A have been implicated in mutagenesis of papillomaviruses and polyomaviruses^3,4^ and, most recently, A3B in restricting and mutagenizing Epstein-Barr virus and Kaposi’s sarcoma herpesvirus (related gamma-herpesviruses)^5^. Although susceptible viruses encode measures to counteract restriction by A3 enzymes, strong cases can be made for each virus leveraging A3 mutagenic potential to promote its own evolution (ex. immune escape). Thus, there is an impetus to develop small molecule inhibitors of A3 enzymatic activity as a means of constraining virus evolution and preventing adverse outcomes such as drug resistance^1,6^).

The antibody gene diversification enzyme AID and several antiviral A3 enzymes have also been implicated in genomic DNA mutations in cancer (reviewed by refs.^7–9^). In particular, A3B is the leading candidate for solid tumor mutagenesis because it is the only constitutively nuclear family member, overexpressed in many different cancer types, active in cancer cell line extracts, and upregulated by cancer-causing viruses (HPV, above). High A3B levels in tumors have also been correlated with poor clinical outcomes, including drug resistance and metastasis^10,11^. A3A and A3H have also been implicated in solid tumor mutagenesis^12–14^, but the endogenous forms of these enzymes have thus far proven challenging to detect and study in model systems such as tumor-derived cell lines.

A striking feature of A3B mutagenesis in cancer is that the resulting mutation spectrum can be reconciled structurally. The APOBEC mutation signature in cancer is defined as C-to-T and C-to-G base substitution mutations within 5′-TC(A/T) trinucleotide motifs (reviewed by refs.^7,8,15,16^). 5′-TCG trinucleotides are also preferred substrates for A3B/A-catalyzed deamination^17–19^ but this motif complicates bioinformatic analyses due to overlap with methyl-CG motifs, which are prone to spontaneous deamination and associate with the biological age of cancer patients (*i.e.* ageing mutation signature). Relative to the target cytosine base (0 position), the 5′ nucleobase (−1 position) is particularly important. In both A3B catalytic domain (A3Bctd*, see below) and A3A x-ray structures with bound ssDNA substrates, the −1 T forms 3 hydrogen bonds with loop 7 residues, 2 direct and 1 water-mediated^19,20^. Other nucleobases, particularly purines, clash with loop 7 residues and are poorly accommodated. Moreover, the integrity of the atomic structures has been validated by systematically changing a single residue (Asp314 in A3Bctd and Asp131 in A3A) to Glu, which alters the hydrogen bonding potential and changes the enzyme’s intrinsic preference from 5′-TC to 5′-CC^19^.

Given the importance of A3B in cancer, small molecule inhibitors of A3B are desirable. However, the lack of A3B-small molecule co-structures constrains such efforts; particularly, structure-guided A3B ligand discovery and optimization. Here we report high-resolution structures of an A3B chimera, one in the presence of a small molecule, as well as dynamics of a catalytically competent A3B construct with an engineered active site, in order to address this knowledge gap and help expedite the rational design of selective A3B inhibitors.

## Results

### Design of A3B-GL7

We previously reported structures of the A3B C-terminal catalytic domain (A3Bctd), determined in several different crystal forms. All DNA-free A3Bctd structures have thus far shown a tightly closed active site, in which loops 1 and 7 on either side of the ssDNA-binding groove associate with each other and block access of ssDNA substrates^21,22^ (Fig. 1). Replacing loop 1 with that from A3A, a Z1-class cytosine deaminase >90% identical to A3Bctd, facilitated structural studies of ssDNA-bound A3B (A3Bctd*)^19^ by freeing-up loop 7 residues for direct interactions with ssDNA substrate, as well as by introducing a histidine into loop1 that helps to hold ssDNA in the active site. We therefore reasoned that, replacing A3Bctd loop 7 with that from A3G catalytic domain, another Z1 cytosine deaminase, may similarly facilitate opening of the active site to allow binding of small molecules. Thus, in this study, we used A3Bctd-QMΔloop3-GL7 (hereafter called A3B-GL7), a highly soluble and catalytically active A3Bctd construct with a grafted loop 7 derived from A3G. Loop 7 residues are critical determinants of the local intrinsic sequence preferences for deamination of each A3 enzyme with, for instance, A3B and A3A preferring 5′-TC substrates and A3G preferring 5′-CC substrates^17,18,23–25^. This intrinsic preference is explained at the atomic level by 2 direct and 1 water-mediated hydrogen bonds between A3B/A loop 7 residues and the −1 T^19,20^ and 3 direct hydrogen-bonds between A3G loop 7 and the −1 C^26^. Consistent with these structural observations, the chimeric protein A3B-GL7 was shown to preferentially deaminate cytosine in a 5′-CC target sequence in DNA, similarly to wild-type A3G^21^.

**Figure 1.**
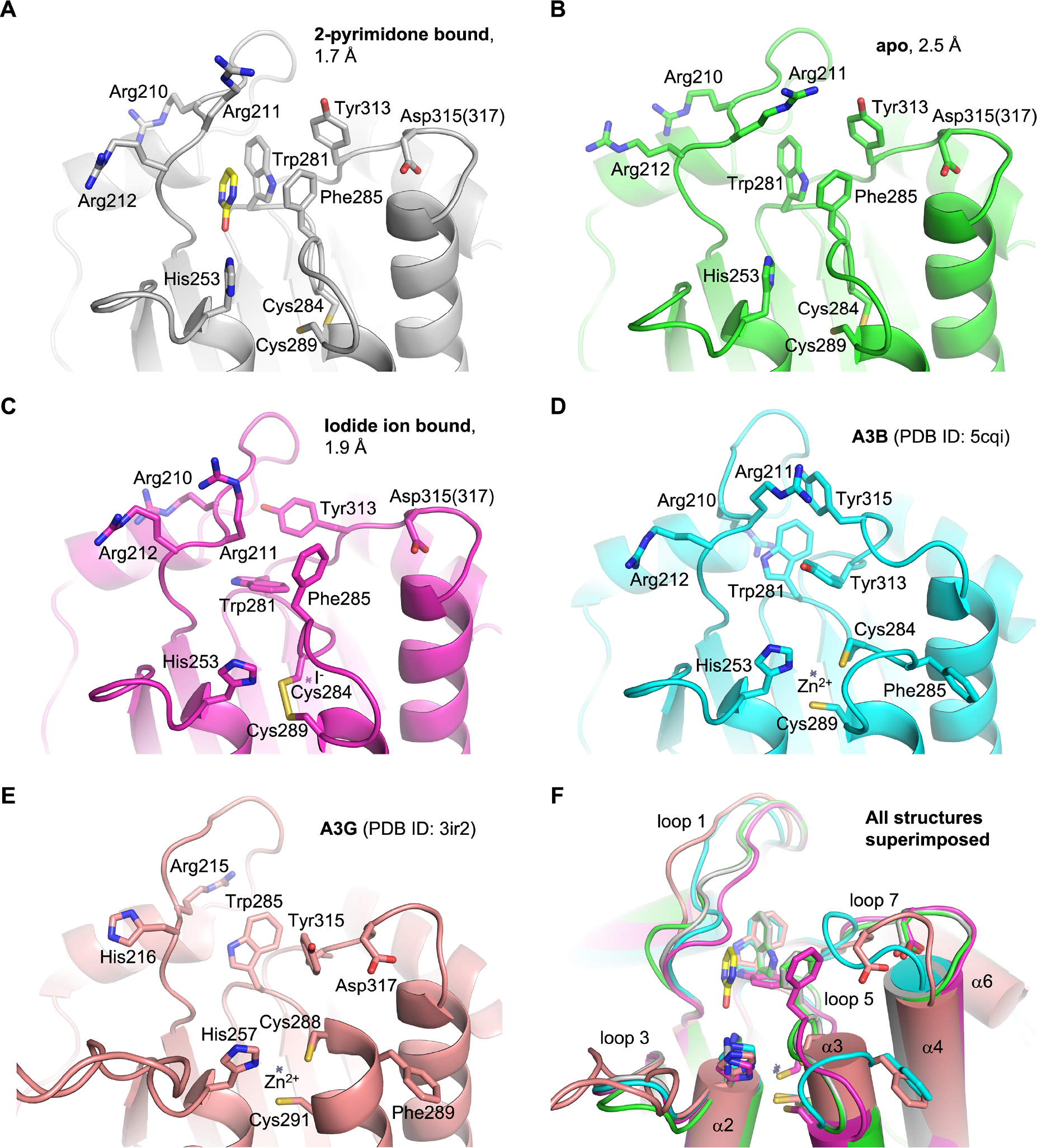
Active site comparisons. Structures around the active site of A3B-GL7 determined in this study (**A**-**C**), compared to those of previously reported A3Bctd structures^21^ (**D**) and A3Gctd structures^43^ (**E**). A superposition of all these structures is shown in (**F**). Key side chains are shown in sticks. Asp315 in A3B-GL7 is labeled according to A3B and corresponds to Asp317 in A3G loop 7. Small crosses mark the positioning of zinc or iodide. 2-pyrimidone bound in the active site of A3B-GL7 (**A**) is shown in sticks (yellow: carbon, red: oxygen, blue: nitrogen).

### A3B-GL7 crystal structure shows a zinc-free active site

We first co-crystallized A3B-GL7 with 2-pyrimidone (2-hydroxypyrimidine), which had been used previously as a mechanism-based inhibitor for zinc-dependent cytosine deaminases^27,28^, and determined its structure at 1.7 Å resolution (Figs. 1A and 2C). The refined structure of A3B-GL7 showed an overall fold very similar to those of prior A3Bctd structures^21,22^ (Figs. 1D and 2A), except that loops 1, 5, and 7 that shape the active site pocket adopt distinct conformations (Fig. 1F). Notably, the electron density clearly indicated that the catalytically essential zinc ion is absent and, accordingly, the zinc-coordinating residues (His253/Cys284/Cys289) have been rearranged. A zinc-free active site was reported previously for a variant of the A3F catalytic domain, where lower pH promoted reversible dissociation of zinc^29^. As our 2-pyrimidone-bound A3B-GL7 crystal was obtained in an acidic condition with pH 5.5, we considered the possibility that loss of the zinc ion was due to low pH or alternatively the very high concentration (0.4 M) of 2-pyrimidone, which may chelate zinc (e.g. PMID: 24997687). Thus, we next crystallized A3B-GL7 in a neutral condition at pH 7.0 in the presence of 10 mM of 2′-deoxyzebularine, a nucleoside analogue bearing 2-pyrimidone as the nucleobase^30^, and determined its structure at 2.5 Å resolution. This structure, unexpectedly, still showed a zinc-free active site similar to that described above, but without a ligand (apo form; Fig. 1B). These results combined to indicate that A3B-GL7 is prone to losing zinc even in a neutral pH condition.

**Figure 2.**
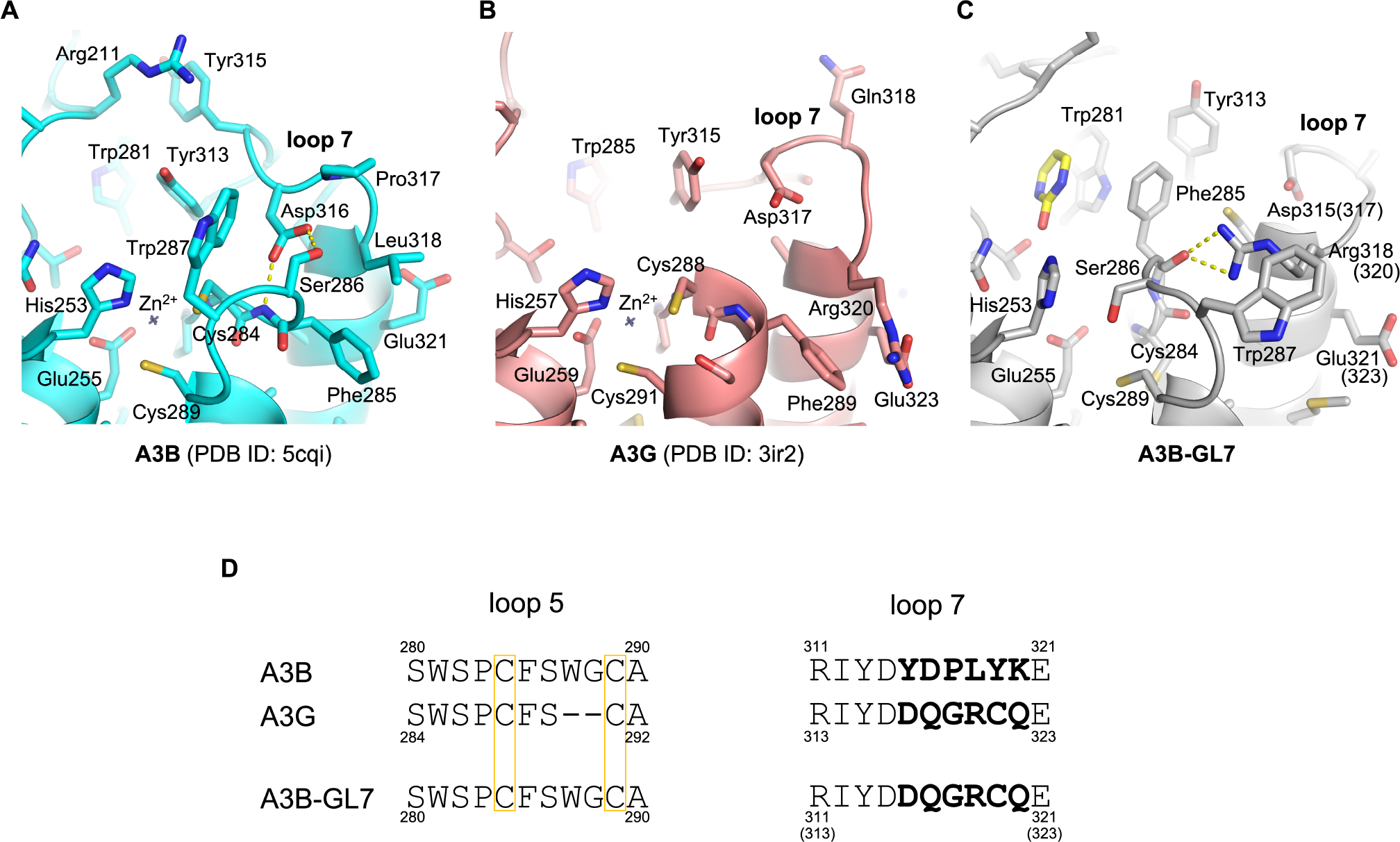
Distinct conformations of loops 5 and 7 from A3B and A3G. Active site conformations as observed in the crystal structures of A3Bctd^21^ (**A**), A3Gctd^43^ (**B**), and A3B-GL7 (**C**), with the side chains of all residues shown in sticks. Key hydrogen bonds between loops 5 and 7 are indicated by yellow dotted lines. (**D**) Amino acid sequence alignment for loops 5 and 7. The zinc-coordinating cysteine residues are boxed. The loop 7 residues that differ between A3B and A3G are in bold.

### Extensive rearrangement observed in the A3B-GL7 active site

The unique conformation of the zinc-free active site of A3B-GL7 features a network of aromatic interactions involving Trp281, Phe285, and Tyr313 (Figs. 1A, B and 2C). Phe285, which is located between the two zinc-coordinating cysteines (Cys284/Cys289) in the amino acid sequence of A3B and is solvent-accessible in the native A3Bctd structures^21,22^, is flipped into the active site. 2-pyrimidone, instead of mimicking the substrate cytosine, fits in a pocket formed between this aromatic cluster and loop 1, being also stabilized by van der Waals contacts with Arg210 backbone and His253 side chain positioned co-planarly with the pyrimidine moiety (Fig. 3A). In this pocket, 2-pyrimidone forms hydrogen bonds with the side chains of Thr214 and Asn240, and the backbone of Gln213 (Fig. 3B), which are positioned similarly in the zinc-bound native A3Bctd active site (e.g. 5cqi; Figs. 1D and 2A). In comparison to the native A3B structure, the loop 1 of A3B-GL7 harboring the stretch of arginines (Arg210/211/212) is shifted inward, slightly toward the active site. In contrast, the extensive rearrangement of the active site residues forces loop 7 away from the active site and closer to the α6 helix (Fig. 1F). Although it is shifted by ~4 Å away from the active site, the grafted loop 7 of A3B-GL7 adopts a similar local conformation as does loop 7 of A3G in its native environment (Figs. 1E, 2B and 4). This suggests structural autonomy of A3G loop 7, which has been demonstrated to play a critical role in determining the target sequence preference^26,31^. The observation helps to explain why A3B-GL7, in its active state, exhibits the A3G-like (5′-CC) target sequence preference^21^.

**Figure 3.**
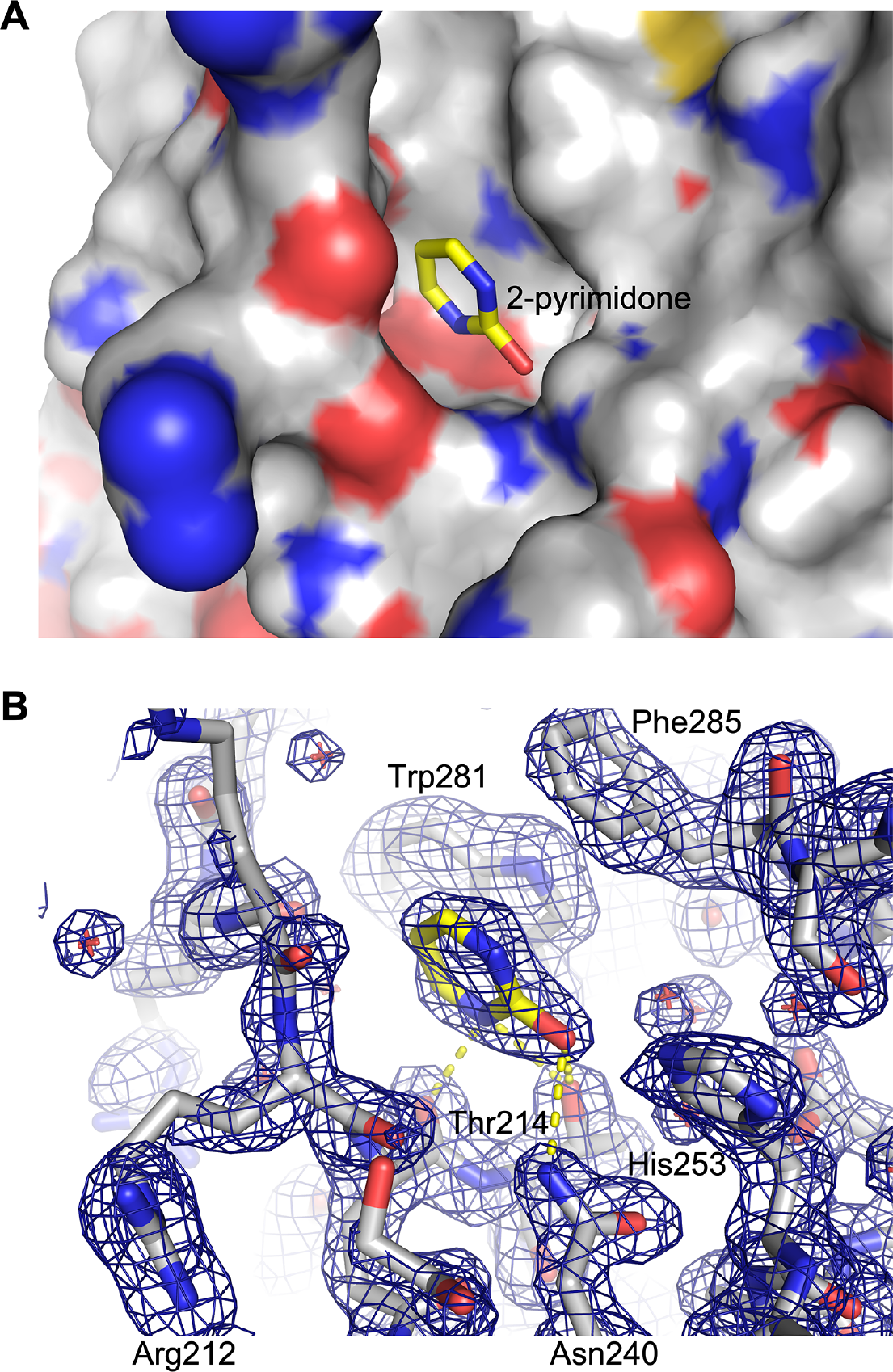
2-pyrimidone bound in the active site. (**A**) Solvent-accessible surface of A3B-GL7 with heteroatoms colored (red: oxygen, blue: nitrogen, sulfur, yellow) and 2-pyrimidone shown in sticks. (**B**) Protein and the bound 2-pyrimidone both in sticks, overlaid with 2mFo-DFc electron density in blue mesh contoured at 1.0 σ. Possible hydrogen-bonds are indicated by yellow dotted lines.

**Figure 4.**
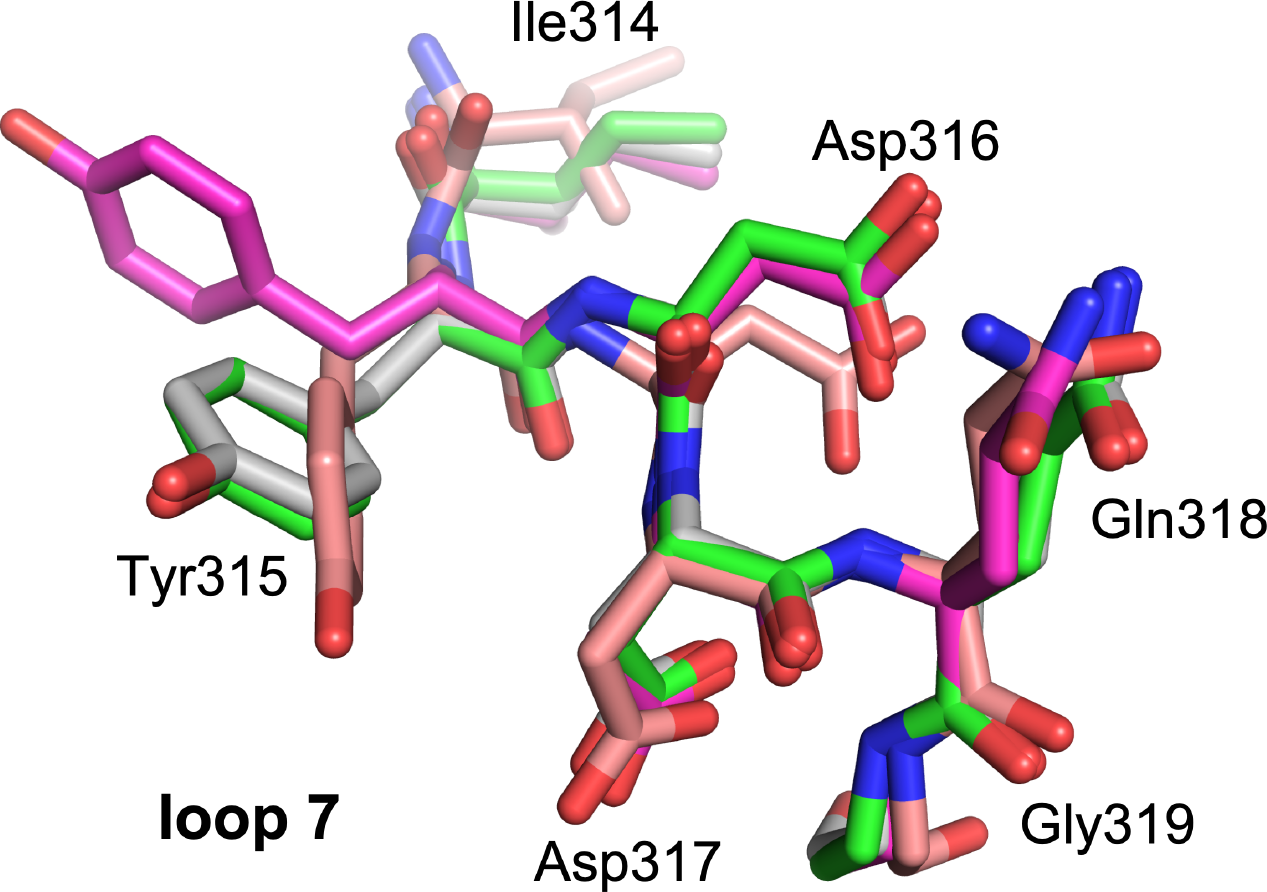
A3G loop 7 conformations. Local superposition of the loop 7 region of A3B-GL7 and A3G structures (Fig. 1A-C and E). Coloring scheme follows that in Fig. 1. The residue numbers are based on to the native A3G sequence.

### Iodide ion stabilizes a *cis* peptide bond in loop 5

In addition to the two structures with and without 2-pyrimidone as described above in similar active site conformations (Figs. 1A, B and 2C), we obtained a structure of A3B-GL7 in a crystallization condition with neutral pH and containing 45 mM sodium iodide (NaI). The resulting structure refined to 1.9 Å resolution again showed a zinc-free active site. However, in this structure loop 5 residues (Trp281-Gly288) are further rearranged and the zinc-coordinating cysteine residues (Cys284/289) formed a disulfide (Fig. 1C). A key feature of this structure is the *cis* conformation for the Ser282-Pro283 peptide bond (Fig. 5A). This is highly unexpected, as these conserved Ser-Pro residues adopt a more thermodynamically favorable *trans* conformation in all other A3 structures reported to date, including those of A3B-GL7 described above (Fig. 5B). In the structure obtained with NaI, we observed a strong (> 15σ) spherical electron density directly adjacent to the flipped proline side chain and surrounded by hydrophobic side chains Trp277, Ile279, and Ile307, which we interpreted to be an iodide ion (Fig. 5A). It is likely that an iodide ion occupying this site stabilized the distorted conformation of loop 5 and facilitated disulfide-bond formation in the zinc-free active site.

**Figure 5.**
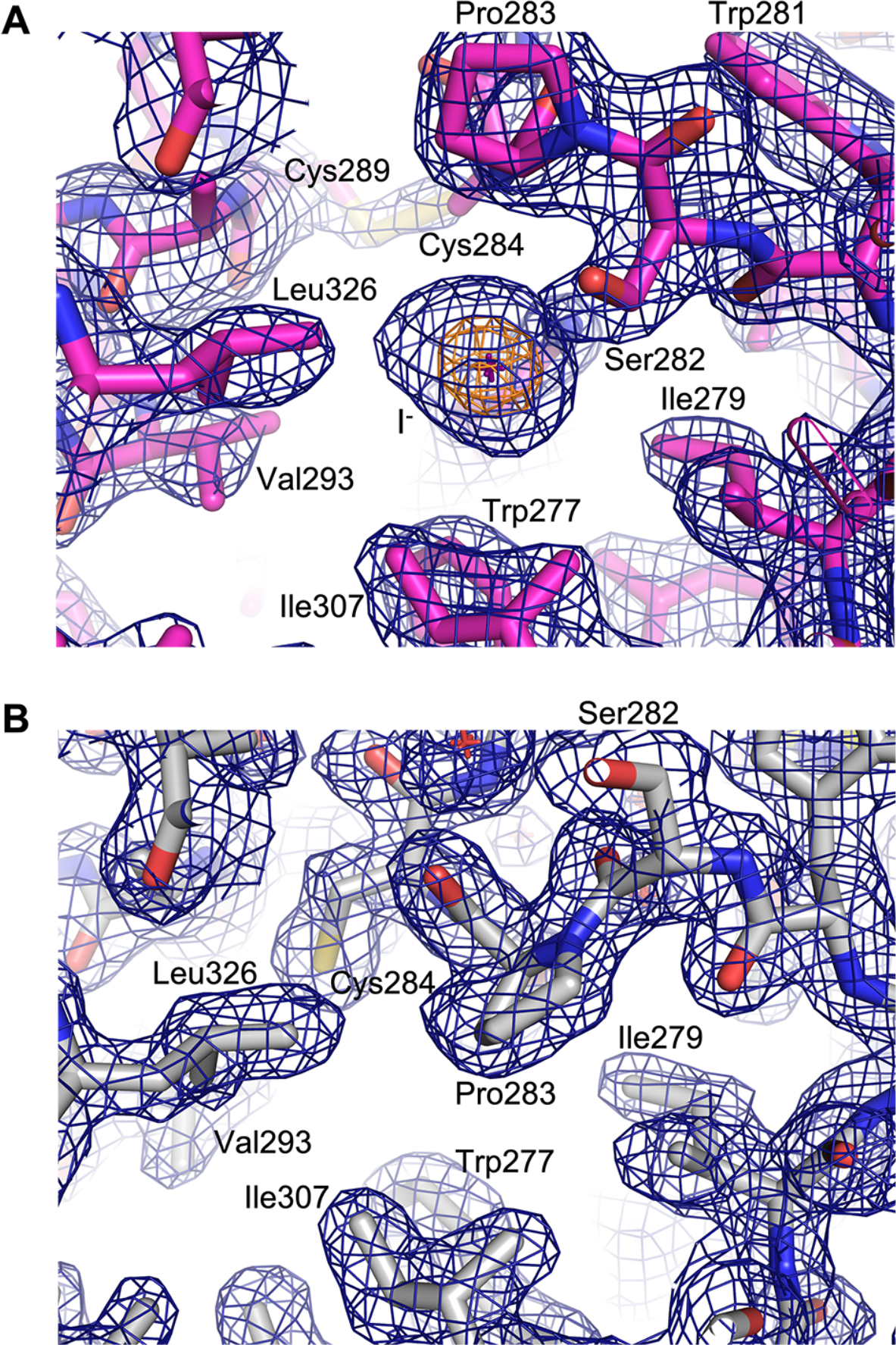
*Cis* vs. *Trans* conformations of the Ser282-Pro283 amide bond. Zinc-less A3B-GL7 structure bound to sodium iodide (**A**, corresponding to Fig. 1C) or with 2-pyrimidone (**B**, corresponding to Fig. 1A) in stick models, overlaid with 2mFo-DFc electron density in blue and orange mesh contoured at 1.0 and 7.5 σ, respectively. The position of iodide ion is marked by a cross. Note that Cys284 is disulfide bonded in (**A**) whereas it is reduced in (**B**), corresponding to *cis* (**A**) and *trans* (**B**) conformations of the adjacent Ser282-Pro283 amide (peptide) bond.

### Molecular dynamics simulations reveal A3B-GL7 active site plasticity

To understand the molecular basis of the extensive rearrangements of A3B-GL7 active site, we performed a series of molecular dynamics (MD) simulations. MD simulations of a computationally constructed zinc-bound A3B_wt_-GL7, which has the wild-type A3B sequence restored otherwise, showed elevated dynamics compared to wild-type A3B, not only for the grafted loop 7 but also the neighboring loops 1 and 5 (Fig. 6A). In wild-type A3B, Asp316 from loop 7 showed persistent hydrogen bonding with Phe285 and Ser286 from loop 5, as observed in the crystal structure (Fig. 2A). These interactions stabilize loop 5 of A3B, which has extra amino acids between the zinc-coordinating Cys residues compared to that of A3G (Fig. 2D), in a unique conformation. In contrast, the grafted loop 7 in A3B-GL7 does not support this native conformation of loop 5. Arg318 from loop 7 (corresponding to A3G Arg320) in the zinc-bound A3B_wt_-GL7 MD simulations predominantly made a cation-π interaction with Phe285 side chain (Fig. 6B), whereas it forms hydrogen bonds with loop 5 backbone in the zinc-free A3B-GL7 crystal structures (Fig. 2C). These observations provide an explanation for the extensive active site rearrangements observed in the crystal structures of A3B-GL7 (Fig. 1A-C), including the loss of the zinc ion.

**Figure 6.**
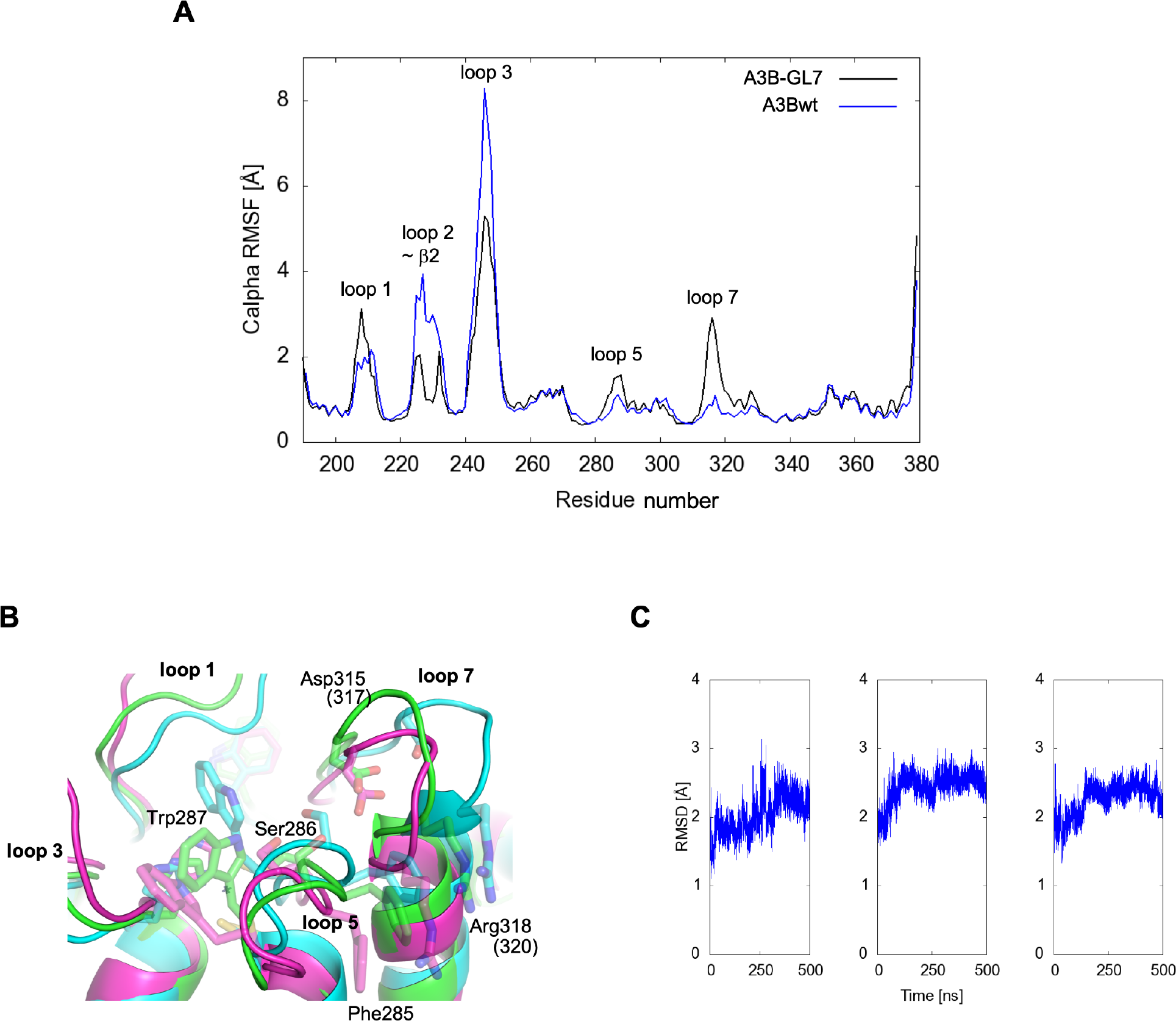
Molecular Dynamics simulations demonstrating active site plasticity. (**A**) RMSF analysis comparing the zinc-bound A3B_wt_-GL7 and wild-type A3Bctd. The A3B_wt_-GL7 starting model was constructed by swapping the loop 7 residues of A3Bctd (similar to Figs. 1D and 2A, but reverted to wild-type computationally) with those from A3G (as in Figs. 1E and 2B). (**B**) Superposition of the final frames of 3 independent 500-ns MD simulations of the zinc-bound A3B_wt_-GL7, showing divergence of the loops that surround the active site. Some side chains are shown in semi-transparent sticks. (**C**) RMSD of Cα atoms with respect to the initial frame for each system plotted to show the stability of MD simulations.

## Discussion

The crystal structures of A3B-GL7 captured in distinct inactive conformations (Figs. 1A-C and 2C), combined with MD simulation results (Fig. 6), highlight remarkable plasticity of the A3B active site and identify key interactions that stabilize the native A3B active site. Of particular interest is the previously unappreciated role of loop 7 of A3B in maintaining the catalytically essential zinc ion, through stabilizing loop 5 in the conformation compatible with zinc-coordination. Although high structural similarity between the A3B and A3G catalytic domains allowed generation of a catalytically competent chimeric enzyme A3B-GL7, this chimeric protein was found to more readily lose the zinc ion than A3B. This is likely to reflect a critical difference between the A3B and A3G active sites; whereas the two zinc-coordinating cysteines of A3G (Cys288/Cys291) are part of an α-helix (Fig. 2B), the corresponding residues Cys284/Cys289 are separated by additional intervening residues as part of loop 5, requiring additional hydrogen-bonding interactions to maintain the conformation compatible with zinc-coordination (Fig. 2A). Typical zinc-finger motifs, in which a set of 4 cysteine or histidine residues coordinate an zinc ion with tetrahedral geometry, are found in numerous proteins and often contribute to protein folding and stability^32–35^. In contrast, the zinc ion in the active site of A3 enzymes is coordinated by 2 cysteines and a histidine, leaving the fourth position of tetrahedral coordination available for engaging the nucleophilic water molecule. Thus, while zinc-binding in the A3B active site is not required for domain folding, it is critically dependent on conformations of the surrounding protein residues.

Our studies also suggest possible means by which small molecules could inhibit A3B activity. Our results show that, while the catalytic zinc ion in APOBEC3 active sites are not essential for the domain fold, loss of the zinc ion causes dramatic rearrangement of the A3B active site. The two distinct inactive conformations that we observed, facilitated by substitution of loop 7 to destabilize the native A3B active site, allowed binding of 2-pyrimidone or an iodide which in turn may have helped stabilize these unique conformations. Even though 2-pyrimidone and iodide ion themselves are not potent inhibitors, they interact with the amino acid residues found in wild-type A3B. We therefore envision that novel small molecules that achieve higher affinity to stabilize these non-productive conformations might serve as chemical inhibitors for A3B, especially given the remarkable conformational plasticity of the A3B active site as demonstrated by MD simulations here, as well as in our prior studies^19,36^. In this regard, the A3B-GL7 structures presented may contribute to the future development of potent A3B inhibitors, which we hypothesize will help to control tumor evolution and other mutational processes relevant to human diseases.

## Materials and methods

### Crystal structure determination

A3Bctd-QMΔloop3-GL7 (A3B-GL7), which has a substitution of A3G residues 317 to 322 (DQGRCQ) for residues 315 to 320 (YDPLYK) of A3Bctd-QMΔloop3, was expressed in *E. coli* strain C41(DE3)pLysS and purified using nickel-affinity and size-exclusion chromatography as reported previously^21^. QM and Δloop3 stand for a set of solubility-enhancing mutations, F200S/W228S/L230K/F308K, and replacement of the loop 3 region spanning Ala242 to Tyr250 by single serine, respectively. The crystal bound to 2-pyrimidone was obtained by the hanging drop vapor diffusion method, by mixing purified A3B-GL7 protein in 20 mM Tris-HCl (pH 7.4), 0.5 M NaCl, 5 mM β-mercaptoethanol and 0.4 M neutralized 2-pyrimidone (2-hydroxypyrimidine)-hydrochloride with a well solution consisting of 0.1 M Bis-Tris-HCl (pH 5.5) and 25% w/v PEG3350. The apo crystal was obtained similarly but in the presence of 10 mM α/β-2′-deoxyzebularine instead of 2-pyrimidone, and a well solution consisting of 0.2 M LiCl, 20% w/v PEG6000, and 0.1 M HEPES-NaOH (pH 7.0). We did not observe electron density for 2′-deoxyzebularine. The iodide-bound crystal was obtained using a E255Q mutant of A3B-GL7 in the presence of a single-stranded oligo DNA containing the CCC target motif, using a well solution consisting of 45 mM NaI and 8.6% w/v PEG3350. The crystal did not contain DNA. All x-ray diffraction data were collected at the Northeastern Collaborative Access Team beamlines (24-ID-C/E) at the Advanced Photon Source, Argonne National Laboratory, and were processed using XDS^37^. The structures were determined by molecular replacement phasing using PHASER^38^, using the A3Bctd-QMΔloop3 structure as the search model. Model building was performed using COOT^39^ and the structures were refined using PHENIX^40^. While we do not observe electron density for hydrogen atoms at 1.7 Å resolution, we chose the 2-pyrimidone tautomer over 2-hydroxypyrimidine to describe the bound ligand in one of the A3B-GL7 structures, based on the hydrogen bonding potentials with surrounding protein groups. The summary of x-ray data collection and model refinement statistics, along with PDB accession codes for the atomic coordinates and structure factors, is shown in Table 1.

**Table 1.**
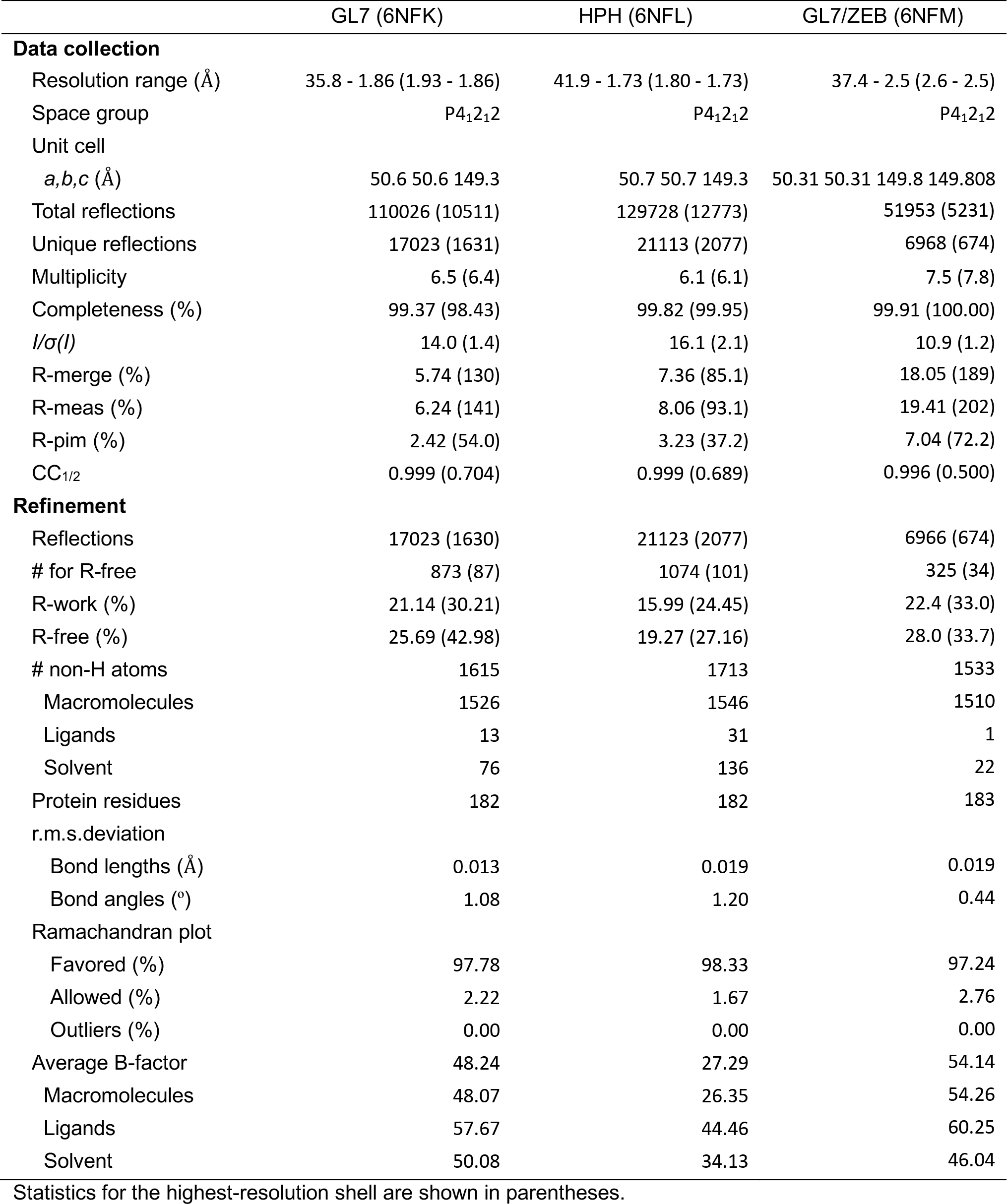
Data collection and model refinement statistics.

|  | GL7 (6NFK) | HPH (6NFL) | GL7/ZEB (6NFM) |
| --- | --- | --- | --- |
| **Data collection** |  |  |  |
| Resolution range (Å) | 35.8 - 1.86 (1.93 - 1.86) | 41.9 - 1.73 (1.80 - 1.73) | 37.4 - 2.5 (2.6 - 2.5) |
| Space group | P4<sub>1</sub>2<sub>1</sub>2 | P4<sub>1</sub>2<sub>1</sub>2 | P4<sub>1</sub>2<sub>1</sub>2 |
| Unit cell |  |  |  |
| *a,b,c* (Å) | 50.6 50.6 149.3 | 50.7 50.7 149.3 | 50.31 50.31 149.8 149.808 |
| Total reflections | 110026 (10511) | 129728 (12773) | 51953 (5231) |
| Unique reflections | 17023 (1631) | 21113 (2077) | 6968 (674) |
| Multiplicity | 6.5 (6.4) | 6.1 (6.1) | 7.5 (7.8) |
| Completeness (%) | 99.37 (98.43) | 99.82 (99.95) | 99.91 (100.00) |
| *I*/$\sigma$(*I*) | 14.0 (1.4) | 16.1 (2.1) | 10.9 (1.2) |
| R-merge (%) | 5.74 (130) | 7.36 (85.1) | 18.05 (189) |
| R-meas (%) | 6.24 (141) | 8.06 (93.1) | 19.41 (202) |
| R-pim (%) | 2.42 (54.0) | 3.23 (37.2) | 7.04 (72.2) |
| CC<sub>1/2</sub> | 0.999 (0.704) | 0.999 (0.689) | 0.996 (0.500) |
| **Refinement** |  |  |  |
| Reflections | 17023 (1630) | 21123 (2077) | 6966 (674) |
| # for R-free | 873 (87) | 1074 (101) | 325 (34) |
| R-work (%) | 21.14 (30.21) | 15.99 (24.45) | 22.4 (33.0) |
| R-free (%) | 25.69 (42.98) | 19.27 (27.16) | 28.0 (33.7) |
| # non-H atoms | 1615 | 1713 | 1533 |
| Macromolecules | 1526 | 1546 | 1510 |
| Ligands | 13 | 31 | 1 |
| Solvent | 76 | 136 | 22 |
| Protein residues | 182 | 182 | 183 |
| r.m.s.deviation |  |  |  |
| Bond lengths (Å) | 0.013 | 0.019 | 0.019 |
| Bond angles (°) | 1.08 | 1.20 | 0.44 |
| Ramachandran plot |  |  |  |
| Favored (%) | 97.78 | 98.33 | 97.24 |
| Allowed (%) | 2.22 | 1.67 | 2.76 |
| Outliers (%) | 0.00 | 0.00 | 0.00 |
| Average B-factor | 48.24 | 27.29 | 54.14 |
| Macromolecules | 48.07 | 26.35 | 54.26 |
| Ligands | 57.67 | 44.46 | 60.25 |
| Solvent | 50.08 | 34.13 | 46.04 |

Statistics for the highest-resolution shell are shown in parentheses.

### Molecular dynamics simulations

Molecular dynamics simulations were performed with Amber18 program^41^ using Amber FF14SB force field^42^ for amino acids, the Cationic Dummy Atom Model (DOI: 10.1007/s008940050119) for the catalytic zinc ion and the zinc-coordinating residues and TIP3P model for water molecules. The starting coordinates of the A3B_wt_-GL7 were generated by modeling in the loop7 of A3G onto the wild-type A3B structure. For the loop7 of A3G to graft, we used residues 313-324 (A3G numbering) of chain A in PDB ID: 3IR2. For the A3B structure, we used the wild-type A3B model we generated from PDB ID: 5CQI by molecular modeling previously^21^. The A3B_wt_-GL7 system was prepared by Leap solvating in a water box with 10 Angstrom buffer, then neutralizing with 7 Na+ ions and finally obtaining a salt concentration of 0.2 M by adding more Na^+^ and Cl^−^ ions. The final system consisted of 35,151 atoms. The system went through multi-step energy minimization, followed by gradual heating and multi-step equilibration with decreasing restraints. The detailed MD protocol used for this system is exactly the same as outlined in our recent publication^36^. The final production MD trajectories with three independent copies, 500 ns each, were generated in an NPT ensemble at 310 K without any restraints. For each MD copy, 10,000 data frames are saved and analyzed. The RMSD plots depicting the stability of MD runs are shown in Fig. 6C.

## Acknowledgments

We thank Stephanie Breunig for synthesizing the 2′-deoxyzebularine and Kayo Orellana for purifying the proteins used in this study. This work was supported by grants from the US National Institutes of Health (NIGMS R01-GM118000 to RSH and HA, NIGMS R35-GM118047 to HA, NIGMS R01-GM110129 to DAH, DP2-OD007237 and NIGMS P41-GM103426 to REA) and the NSF (CHE060073N to REA). This work is based upon research conducted at the Northeastern Collaborative Access Team beamlines, which are funded by the US National Institutes of Health (NIGMS P30 GM124165). The Pilatus 6M detector on 24-ID-C beamline is funded by a NIH-ORIP HEI grant (S10 RR029205). This research used resources of the Advanced Photon Source, a U.S. Department of Energy (DOE) Office of Science User Facility operated for the DOE Office of Science by Argonne National Laboratory under Contract No. DE-AC02-06CH11357, and those of the Minnesota Supercomputing Institute. R.S.H. is the Margaret Harvey Schering Land Grant Chair for Cancer Research, a Distinguished McKnight University Professor, and an Investigator of the Howard Hughes Medical Institute.

